# Cross-linker binding strength influences actin assembly microstructure and myosin II driven remodeling

**DOI:** 10.64898/2026.10.02.756345

**Authors:** Carolyn A. Feigeles, Megan Janquart, Veronica Armstrong, Kimberly L. Weirich

## Abstract

The actin cytoskeleton is a self-assembled material that acts as the mechanical scaffold of the cell. Within the cell, protein cross-linkers organize actin filaments into assemblies with various microstructures that support distinct physiological functions. These protein cross-linkers differ in structural features. To decouple the role of individual cross-linker properties in actin assembly microstructure and mechanics, we synthesize programmable cross-linkers using a DNA linker coupled to binding domains. We find that high affinity cross-linkers induce actin bundling, while low affinity cross-linkers form actin networks. Myosin II motors remodel and contract all cross- linked networks tested in distinct ways from uncross-linked networks. We find that networks with high affinity cross-linkers have different myosin induced dynamics than networks with low affinity cross-linkers. This research expands our understanding of how cells regulate their mechanics and microstructures, while also providing insight towards the structure-property relationship in complex soft materials.

## Introduction

The cytoskeleton is a dense mixture of self-assembled biopolymer filaments and associated proteins responsible for cellular functions such as regulation of mechanics and transport^1^. The assembly of these cytoskeletal filaments, such as actin, microtubules, and intermediate filaments, is mediated by protein cross-linkers, but has also been shown to result from interactions induced by motor proteins, biomolecular condensates, and through depletion-attraction effects^2–11^. Within the cell, actin forms various cross-linked microstructures, including networks, bundles, and networks of bundles, which enable distinct physiological functions^2,12^. For example, short and rigid cross-linkers, like fascin, constrain actin filaments into tight, parallel bundles which are more rigid than individual actin filaments^13,14^. These stiff microstructures enable cellular protrusions, like filopodia, to withstand compressive forces from their surrounding environment^15,16^. In contrast, long and flexible cross-linker, such as filamin, mediate the formation of actin networks that comprise the cellular cortex^17,18^. Mutations or overexpression of cross-linkers may impact cytoskeletal functions and have been associated with disease^19–21^.

Many physiological cross-linkers share a similar structure, consisting of a linker region as well as two or more actin-binding domains^2,22^. While physiological cross-linkers share basic structure, properties such as their length, rigidity, and binding affinity, can differ^2^. Physiological cross-linkers are often different in more than one property, and the biological toolkit of cross-linkers represents a small collection of the potential cross-linker space, making it challenging to experimentally investigate the effect of individual cross-linker properties on actin network microstructures and mechanics. Systematic investigations of this structure-property relationship could improve our understanding of cytoskeletal mechanics, as well as inform computational models. One method that has been used to overcome these challenges is to create a synthetic cross-linker with tunable properties. In previous work, synthetic DNA cross-linkers have been used to model physiological cytoskeletal networks using networks of peptides and actin filaments^23,24^. Similarly, actin cross- linkers have also been synthesized using a polyethylene glycol linker backbone with LifeAct acting as the actin binding domain^25^. These synthetic cross-linkers provide useful groundwork to study the influence of cross-linker properties on cytoskeletal assemblies.

Cross-linkers have also been known to influence motor driven activity and remodeling of cytoskeletal networks. Motor proteins have been shown to preferentially localize to particular bundle microstructures^26^. Features of motor transport, such as run length, stepping and transport dynamics, as well as myosin forces are also dependent on bundle structure^10,27–29^. Cross-linkers can also impact motor driven contractility through connectivity, which influences the contractile domain size, filament organization, and microscopic mechanisms of deformation, such as filament sliding or buckling^30–37^.

Here, we synthesize a programmable cross-linker modified from Lorenz et al. with variable length and binding affinity to investigate the role of cross-linker properties on actin microstructure and contractile behavior^24^. Using confocal microscopy, we show that high affinity cross-linkers form bundled networks of actin filaments, while low affinity cross-linkers form actin networks. Myosin II motors remodel and contract cross-linked networks, regardless of cross-linker binding affinity, with different dynamics than uncross-linked networks. Interestingly, cross-linker length had less of an effect than cross-linker binding affinity on microstructure and myosin activity.

## Materials and Methods

### Protein Preparation

Proteins were either purified or prepared from commercially available lyophilized powder, then flash frozen in liquid nitrogen, and stored at -80°C until use. Concentration of proteins were determined spectrophotometrically (Fisher Scientific, NanoDrop 2000c Spectrophotometer) using an extinction coefficient for actin of 42680 M^-1^cm^-1^ at 280 nm and ∼148,000 M^-1^cm^-1^ at 280 nm for skeletal muscle myosin II^38^. To determine the ratio between labeled and unlabeled protein, extinction coefficients of ∼90000 M^-1^cm^-1^ at 545 nm for tetramethylrhodamine and 270,000 M^-1^cm^- 1^ at 651 nm for Alexa 647 were used, as provided in the product specifications.

Fluorescently labeled monomeric actin was purified from skeletal muscle acetone powder (rabbit, Pel-Freez Biologicals) according to Spudich et al., labeled with tetramethylrhodamine-6- maleimide (TMR actin), and stored in 2 mM Tris, 0.1 mM CaCl_2_, 0.5 mM DTT, 0.2 mM ATP, 1 mM NaN_3_, pH 8^39^. Unlabeled actin protein (>99% pure, rabbit skeletal muscle, Cytoskeleton), Biotinylated actin (99% pure, rabbit skeletal muscle, Cytoskeleton), and TMR actin (Rhodamine actin protein, >99% pure from rabbit skeletal muscle, Cytoskeleton) were prepared from lyophilized powder by hydrating to 10 mg/mL and pipette mixing over ice for 15 minutes. Then, the actin solution was dialyzed against 2 mM Tris, 0.1 mM CaCl_2_, 0.5 mM DTT, 0.2 mM ATP, 1 mM NaN_3_, pH 8, for at least three hours while stirring over ice. The solution was further dialyzed overnight at 4°C without stirring.

Skeletal muscle myosin II was purified from skeletal muscle (chicken, Bethel Trails Farm, Gray Court, SC) following a protocol derived from the literature^40^. To allow for fluorescent imaging, the myosin monomers were labeled with Alexa-647 maleimide (Invitrogen) using click chemistry following methods previously described^38^. The myosin was stored in 600 mM KCl, 10 mM EDTA, and 25 mM KPO_4_ at pH 6.6 at -80°C.

### DNA Cross-linker Synthesis

The following DNA sequences were used. Each sequence is represented starting from the 5’ end.

#### 20 nm (60 base pair) Cross-linker Sequence

F: GCTTTCTTCTCTAAATACATCTTCACGTCGATATCACCATAACTCAGGTAAGGAGGTCAA R: TTGACCTCCTTACCTGAGTTATGGTGATATCGACGTGAAGATGTATTTAGAGAAGAAAGC

#### 7 nm (20 base pair) Cross-linker Sequence

F: TTGACCCTGTTAGAGAAAGC

R: GCTTTCTCTAACAGGGTCAA

#### High Affinity Cross-linker

To prepare high binding affinity cross-linkers, biotin-streptavidin bonds were used by bridging biotinylated actin and biotinylated DNA with streptavidin (New England Biolabs). Biotin functionalized (5’) DNA oligos (IDT Technologies) were purchased with standard desalting and used without further purification. Oligos were resuspended from lyophilized powder to 100 µM by adding milliQ water. Complementary strands (F and R for each length) were melted by heating to 95°C and held for 10 minutes, transferred to 72.5°C and held for 15 minutes, and quenched in ice for 2 minutes to hybridize strands. Following hybridization, the solution was stored at 4°C until use.

#### Low affinity Cross-linker

To synthesize DNA cross-linkers with phalloidin binding moieties, we adapted the protocol established by Lorenz et al.^24^. Amine-functionalized (5’) DNA oligos were purchased with standard desalting (IDT Technologies). The DNA oligos were hydrated from lyophilized powder to 100 µM in milliQ water and stored at 4°C until use. Stocks of phalloidin amine (AAT Bioquest) and 6- Azidohexanoic Acid (Sigma-Aldrich) were resuspended in dry dimethyl sulfoxide (DMSO) at 5 mM, and Dibenzocyclooctyne-N-hydroxysuccinimidyl (DBCO-NHS) (Sigma-Aldrich) was resuspended in dry DMSO at 10 mM and stored at -20°C until use.

Amino-DNA oligos were added to a concentration of 5 µM for each of set of complementary oligos (F and R) in 100 mM potassium phosphate buffer (20.8 mM KH_2_PO_4_ and 79.2 mM K_2_HPO_4_). The oligo solution was then heated to 95°C for 10 minutes, transferred to 72.5°C, held for 15 minutes, and finally quenched by placing on ice for 2 minutes to hybridize the oligos. Immediately following hybridization, the resuspended DBCO-NHS ester was added to a final concentration of 1 mM in the oligo solution. Separately, a solution of 5 mM phalloidin amine and 5 mM 6-azidohexanoic acid was prepared in 75 mM potassium phosphate buffer (5.2 mM KH_2_PO_4_ and 59.4 K_2_HPO_4_). Both solutions were incubated overnight in dark, ambient conditions.

Then the DNA was isolated using ethanol precipitation. First, sodium acetate was added to the DNA solution to a final concentration of 0.6 M. Cold pure ethanol (200 proof, -20°C) was added to the solution at a volumetric ratio of 3:1 and incubated for a minimum of 2 hours at -80°C. The solution was then centrifuged at 16,000 rcf for 30 minutes at 4°C, resulting in a pellet of DNA. The pellet was rinsed with 75 vol% ethanol three times and air dried for a minimum of 30 minutes. Once dry, the pellet was resuspended in 10 mM Imidazole, 50 mM KCl, 1 mM CaCl_2_, and 0.2 mM ethylene glycol-bis(β-aminoethyl ether)-N,N,N′,N′-tetraacetic acid (EGTA), pH 7.5. Once fully resuspended, the DNA and phalloidin solutions were mixed equivolume and allowed to incubate overnight in dark, ambient conditions.

To remove unreacted phalloidins, the solution was filtered by centrifuging at 14,000 rcf through a 10 kDa filter (Amicon, Sigma-Aldrich) for 5 minutes. The presence of DNA cross-linker in the retentate was confirmed spectrophotometrically (Fisher Scientific, NanoDrop 2000c Spectrophotometer) at 260 nm using extinction coefficients provided by the manufacturer (IDT Technologies). The final solution was stored at 4°C for up to 3 months before use.

#### Sample Preparation for fluorescence microscopy

The sample chamber was composed of a borosilicate cloning cylinder (Corning) epoxied to the center or a borosilicate coverslip (Fisherbrand). Both components were rinsed with ethanol and dried using filtered air. Immediately before adding the sample, the coverslip surface was passivated using 2 wt% 008-Fluorosurfactant in HFE7500 oil and excess oil was removed via pipette.

##### Low affinity cross-linked networks

To prepare networks, 2.64 μM actin monomer (0.264 µM labelled with TMR) was polymerized in a buffer solution containing 10 mM Imidazole, 50 mM KCl, 1 mM CaCl_2_, and 0.2 mM ethylene glycol-bis(β-aminoethyl ether)-N,N,N′,N′-tetraacetic acid (EGTA), pH 7.5, 0.50 wt% *β*- mercaptoethanol, 4.5 mg/mL glucose, 2.7 mg/mL glucose oxidase, 1700 units/mL catalase, 1 mM ATP, 0.4 wt% methylcellulose. DNA cross-linkers were added at concentrations between 2.5 to 20 mol% crosslinker:actin monomer to the polymerized actin network and incubated for 10 minutes to ensure the cross-linker has enough time to cross-link the actin filaments. To initiate network remodeling and contraction, myosin II motors were added to a concentration of 0.125 µM or 0.245 µM to induce coarsening of the cross-linked networks.

##### High affinity cross-linked networks

To prepare actin networks for biotinylated cross-linkers, actin monomer (0.264 µM labelled with TMR, 0.132 µM functionalized with biotin) was polymerized by adding to the buffer described in the above sample. The biotinylated actin concentration was chosen to minimize bundling between actin filaments and streptavidin (from a 1mg/mL stock in water, New England Biolabs), while also providing enough active binding sites for the cross-linker to interact (See *Supporting Information*). Biotinylated DNA cross-linkers were mixed with streptavidin at a 1:4 molar ratio (cross-linker to streptavidin) and incubated while the actin filaments polymerized. The cross-linker solution was added to the biotinylated actin network and incubated for 10 min before adding myosin II motors (0.125 µM) to induce network remodeling.

#### Confocal Microscopy

Fluorescence samples were imaged using a Nikon Eclipse TI2 microscope equipped with a W1 spinning disk confocal head (Yokogawa) via an ORCA-Fusion CMOS camera (Hamamatsu) and a 60× Apo TIRF objective, 1.49 numerical aperture (Nikon). The samples were illuminated using 561 and 640 nm laser lines.

#### Transmission Electron Microscopy

Actin samples were deposited on 200 mesh copper grids with 5-6 nm carbon support films (Electron Microscopy Sciences) and allowed to incubate for 3-5 minutes before wicking to remove excess solution. The samples were negative stained by incubating with a drop of 1% (w/v) uranyl acetate for 1 min, rinsing in DI H_2_O, and wicking off the remaining solution. Each sample was allowed to air dry for 3 minutes before being placed in a desiccated environment until use. Transmission electron microscopy samples were imaged using a HT7830 UHR (Hitachi) at 120kV accelerating voltage.

#### Image Analysis

To determine the bundle radius, a line was drawn perpendicular across the bundle using ImageJ, with sufficient length to include approximately 1 µm of background on either side of the bundle. When possible, bundles were selected to minimize interference with other nearby actin filaments. ImageJ’s built-in plot profile tool was used to gather the intensity as a function of distance along the line. Each bundle was measured at 3 places, and 40 different bundles were measured for each condition. The intensity fit to the following function:

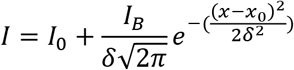

where *I* is the measured intensity, *I*_0_ is the background intensity, *I*_B_ is the bundle intensity, δ is the standard deviation of the intensity distribution, *x* is the distance along the line, and x_0_ is the distance to the maximum intensity from the start of the line. Measurements for the 7 nm biotin crosslinker gaussian fit had an average R^2^ = 0.972 and a minimum R^2^ = 0.847, while the 20 nm biotin crosslinker measurements had an average R^2^ = 0.984 and minimum R^2^ = 0.944. Using the standard deviation from the Gaussian fit, the radius of the bundle, *r*, was calculated from the full width half max of the Gaussian, 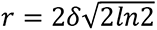.

For myosin cluster analysis, a systematic way to determine a starting frame for the analysis was necessary, since myosin accumulates in the networks at different times. To determine the starting frame, a binary threshold was applied to the myosin channel of each image sequence using ImageJ and the analyze particles function was used to collect the size of each myosin cluster in thresholded frames. To analyze merge events, the frame in which 30% of myosin clusters had an area greater than 0.7 μm^2^ was chosen to be the analysis starting point. Each myosin cluster was tracked, and the number of merging events were noted from the starting point for 45 frames (132 seconds, corresponding to a period of network remodeling) for each movie.

To determine the mean distance between myosin clusters, ImageJ was used to apply a binary threshold to the myosin channel of each image sequence. Once binarized, ImageJ’s Find Maxima feature was used to select each individual myosin cluster. Using a homemade macro, ImageJ’s

Delaunay Voronoi plugin was automated to calculate the number of puncta and the mean distance between clusters over time.

## Results and Discussion

### High affinity, long cross-linkers induce bundling

We synthesized a programmable cross-linker inspired by a cross-linker reported by Lorenz et al., consisting of a double stranded DNA linker region that is approximately 20 nm long (60 base pairs) with binding sites on either end (See *Methods*)^24^. We chose to make a high affinity cross- linker, using biotin-streptavidin to bond the cross-linker and actin filaments together. Biotin and streptavidin are known to form highly specific and stable, high affinity interactions, resulting from the four biotin binding sites per streptavidin molecule^41,42^. Biotin-streptavidin interactions have been used in a multitude of applications, including targeted drug therapies and single molecule protein studies^43–45^. In the high affinity cross-linker, each single stranded DNA oligo has a biotin functionalization at the 5’ end, resulting in biotin at either end of a hybridized pair of complementary oligos. Prior to adding the cross-linker to an entangled actin network, the cross- linker is incubated in a streptavidin solution such that the molar ratio is 1:4 cross-linker:streptavidin (See *Supporting Information*).

We polymerize actin monomers into filaments to form entangled actin networks (Fig. 1A i). The actin filaments have a small fraction of biotinylated actin monomers (5 mol%), such that biotin DNA-streptavidin-biotin actin interactions can form, resulting in a cross-linker bound to an actin filament. To form the cross-linked networks, we add a 20 nm high affinity cross-linker, which mimics the basic structure of physiological cross-linkers, such as alpha-actinin (Fig. 1B i).

**Figure 1:**
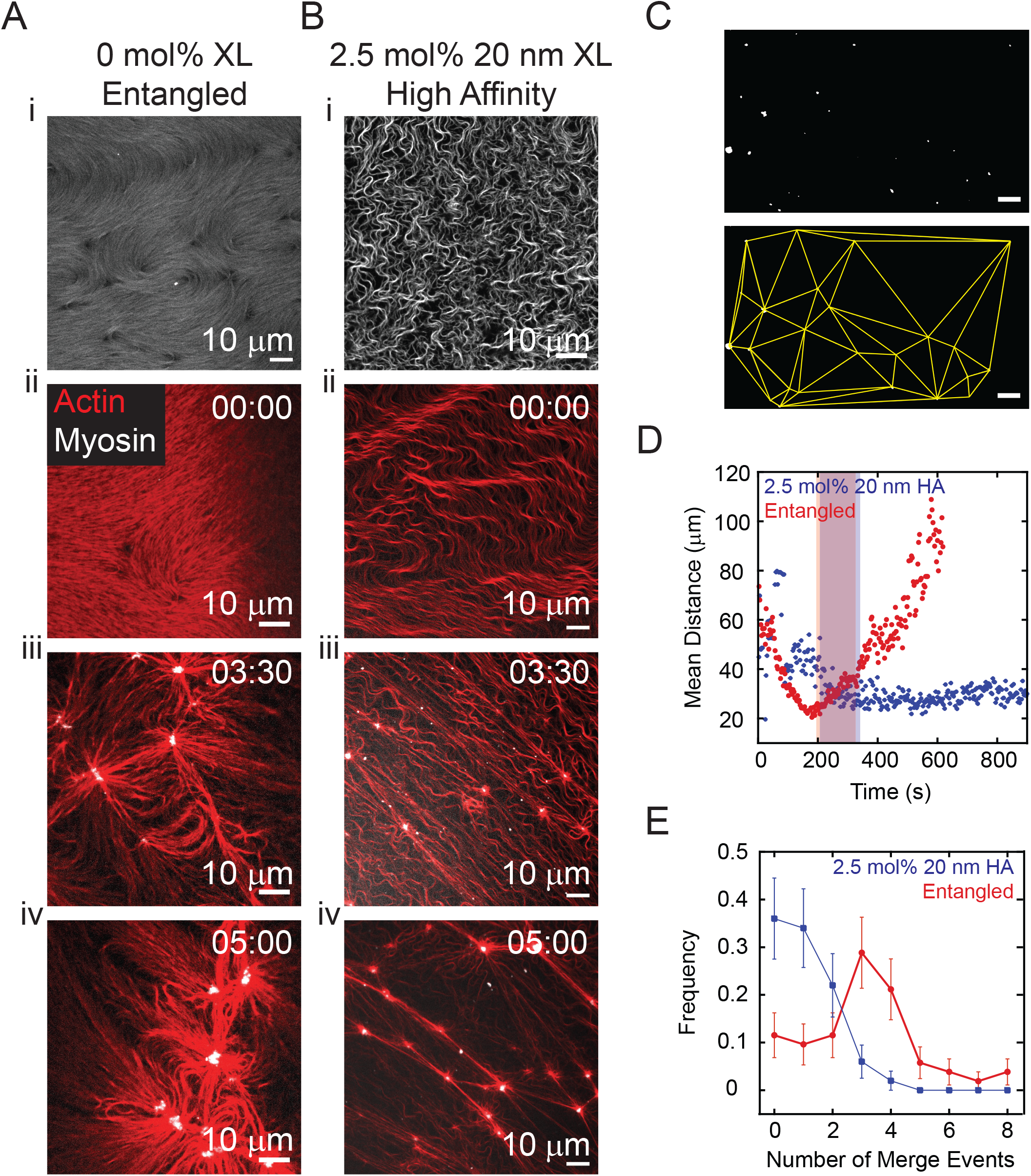
Programmable cross-linker bundles actin filaments. (A) i. Fluorescence microscopy image of 2.64 µM entangled actin filaments. ii-iv. Fluorescence microscopy image sequence of 2.64 µM entangled actin filaments (red) with 0.125 µM myosin II motors (white) added at *t* = 0 s. (B) i. Fluorescence microscopy image of 2.64 µM actin cross-linked with 2.5 mol% 20 nm high affinity cross-linker. ii-iv. Fluorescence microscopy image sequence of an actin network (red) cross-linked by 2.5 mol% 20 nm high affinity cross-linker with 1.25 µM myosin II motors (white) added at *t* = 0 s. (C) Top: Binarized image of myosin II clusters (white). Same data as Fig. 1B iii. Bottom: Delaunay triangulation (yellow) corresponding to myosin image above. (D) Mean distance between myosin clusters over time in entangled (red) and cross-linked (blue) networks, corresponding to data in Fig. 1A and B. (E) Frequency of the number of times a myosin cluster merges within entangled (red) and cross-linked networks (blue). Shaded regions in part D indicate the window of time used to track myosin merge events. Scale bar is 10 µm and time is represented in min:s in all images.

Upon adding the programmable cross-linker to the entangled actin network at a ratio of 2.5 mol% (2.5 mol cross-linker for every 100 mol actin monomers), the actin filaments form an interconnected network of bundles (Fig. 1B i). It is well known that entangled actin exhibits similar bundle formation in the presence of various physiological cross-linkers, such as α-actinin, fimbrin, and fascin^2^. To quantify the bundling, we use the fluorescence intensity as a measure of the amount of actin in a bundle (Fig. S2). To extract the bundle diameter, we measure the intensity across a line that is drawn perpendicular to the length of a bundle and fit the intensity to a Gaussian. From the full width at the half maximum, we find that the bundles had an average diameter of approximately 0.57 ± 0.125 µm. Notably, actin filaments are polar polymers^46^. Physiological cross-linkers are known to form different types of bundles, such as unipolar, where filaments are arranged with the polarity aligned, and mixed polarity bundles^12^. Given the four binding sites for streptavidin to bind to biotin, we expect this cross-linker to only form mixed polarity bundles.

### Myosin II contracts high affinity, long cross-linked networks

To investigate myosin II-induced contractility, we introduced myosin II motors to the cross-linked networks. Upon adding myosin (white), we observe mass movement of the actin network (red), likely induced by myosin in areas outside of the field of view (Fig. 1B ii-iv). Small puncta of myosin (white) appear within the network, as actin bundles buckle (*t*<2:30 min). Myosin puncta aggregate into clusters, and actin bundles between myosin clusters straighten as the myosin clusters move closer together, indicating that the bundles are under tension (Fig. 1B iii), consistent with a contractile cross-linked network.

To quantify the apparent myosin collective movement, Delaunay triangulations (Fig. 1C, bottom, yellow lines) were used to identify a set of nearest neighbors for each myosin cluster (Fig. 1C, top, white)^47,48^. From the length of the sides of the triangles, we calculate an average distance between clusters. During initial times, there is a transient increase in the average distance as myosin puncta form in the networks (Fig. 1D, blue). Then, the mean distance decreases as myosin II approach each other and merge into larger clusters (Fig. 1D, blue, *t* < 200 s). Finally, at later times, the mean distance asymptotes to a steady separation value, as the network begins to move collectively, with myosin clusters remaining in approximately the same configuration with actin bundles extended between them (Fig. 1D, blue, *t* > 200 s). This is consistent with previous studies that show that cross-linked networks can get arrested in a steady state^36,49^.

By contrast, when myosin II motors are added to an entangled network without cross-linkers, we observe different network and myosin coarsening dynamics (Fig.1Aii-iv). Initially, the myosin form puncta and begin clustering while the actin network restructures, with increasing variation in filament packing as well as filament bending (Fig. 1A ii). Once the myosin motors form clusters, actin filaments form small asters interconnected by regions of straight actin filaments (Fig. 1A iii). Finally, the clusters merge to form large, isolated asters that contain most of the actin filaments (Fig. 1A iv).

Again quantifying this with Delaunay triangulation, we find that the mean myosin cluster spacing initially decreases as myosin clusters accumulate within the entangled network and move closer to each other (Fig. 1D, red, *t*<200 sec), similarly to the cross-linked network. As myosin clusters merge to form larger clusters, the mean distance increases (Fig. 1D, red, *t*>200 s), in contrast to the cross-linked network where the myosin spacing stabilizes at long times.

We also tracked the number of times individual myosin clusters within a network merged with neighbors to quantify the differences in clustering dynamics between the cross-linked and entangled networks. Since myosin clustering times vary by network type, we focused on a range of times after myosin clusters had reached a requisite size (See *Methods*). On average, in entangled networks, myosin clusters merged 3.7 ± 1.9 times (n=52), with 20 clusters merging more than four times and two merging up to eight times (Fig. 1E, red). By contrast, in cross-linked networks, myosin clusters merged fewer times, with an average of 1.0 ± 0.68 merges (n=50), and zero clusters that merged more than five times, consistent with a network that approaches an arrested configuration (Fig. 1E, blue).

### Low affinity cross-linkers form actin networks

While the high affinity cross-linker can bundle actin filaments, it represents a much stronger binding interaction than physiological actin cross-linkers^2,41^. Physiological actin cross-linkers bind and unbind the actin filaments within the cross-linked network^2^. For example, physiological cross- linkers such as α-actinin and filamin have equilibrium dissociation constants of 10^-7^-10^-6^ M^50,51^. This transient cross-linking is key to actin network dynamics and re-organization^52,53^. To investigate the effect of transient binding, and decrease the strength of interaction between the filaments and cross-linker, we modified the cross-linker design to replace the biotin with phalloidin, a toxin known to bind to actin filaments^54,55^. Since biotin has a K_d_ of 4.0x10^-13^ M to streptavidin, and phalloidin has a K_d_ of 3.6x10^-8^ M to actin, this change represents five orders of magnitude increase in the equilibrium dissociation constant^56,55^.

When we add 2.5 mol% of the low affinity cross-linker to actin networks, we do not see visible bundling of actin filaments or evidence of network remodeling (Fig. 2A i), in contrast to the high affinity cross-linkers (Fig. 1B). Notably, while physiological cross-linker binding affinity is low and transient, when at a high enough concentration compared to the actin filaments, cross-linkers can form actin bundles^22^. We next increased the concentration to 40 mol%, and we find the filaments in the networks remain unbundled (20 mol% Fig. 2B i; for other concentrations see Fig. S3). As the concentration of cross-linker increases, the actin has visible changes. These changes in the actin network potentially result from phalloidin increasing the stiffness of actin filaments in addition to blocking depolymerization of filaments^54,57^.

**Figure 2:**
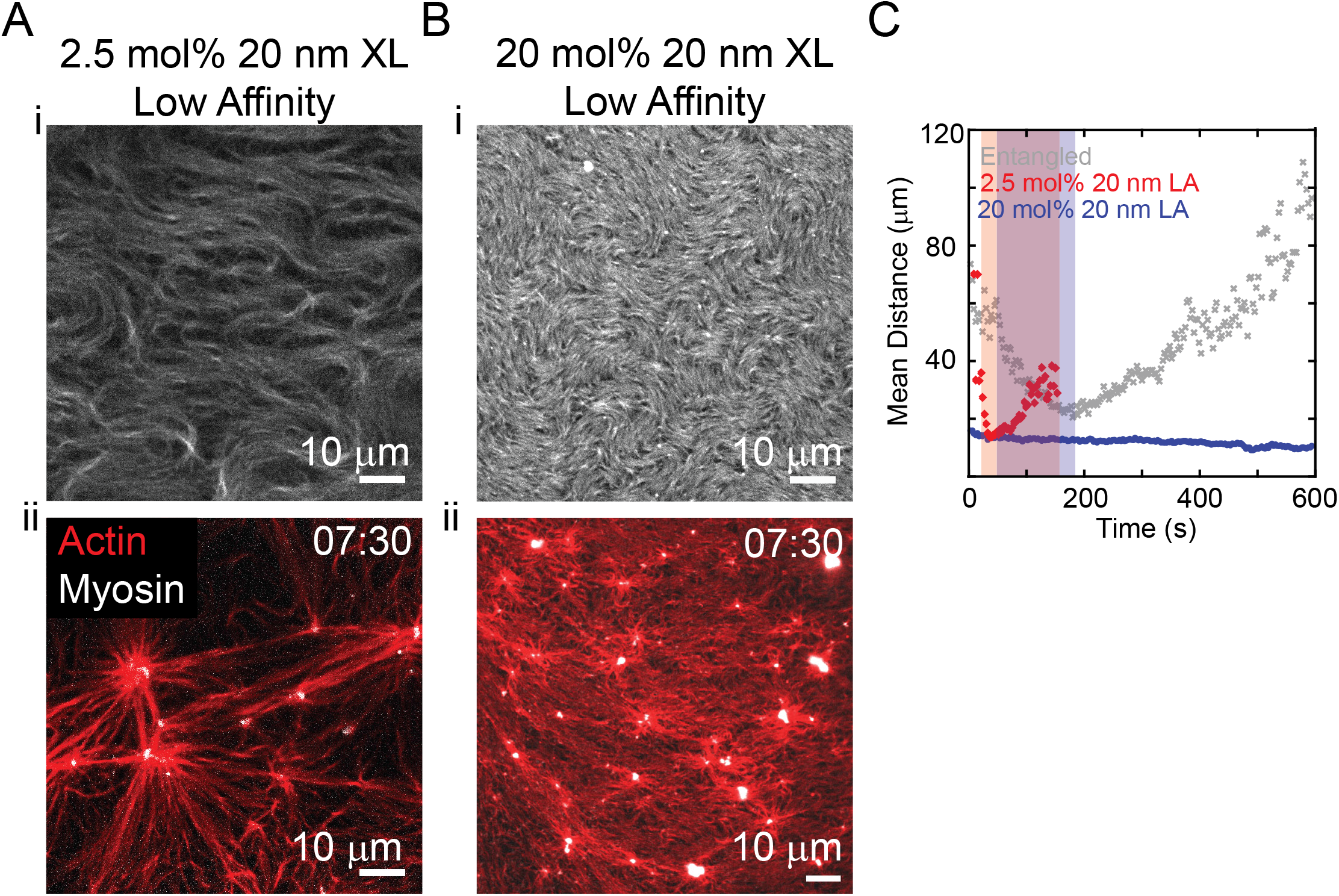
Low affinity cross-linker forms actin networks. (A) i. Fluorescence microscopy images of 2.64 µM actin cross-linked by 2.5 mol% 20 nm low affinity cross-linker. ii. Fluorescence microscopy image of 2.5 mol% 20 nm low affinity cross-linked network (red) 7 min 30 s after 0.12 µM myosin II motors (white) were added. (B) i. Fluorescence microscopy image of 2.64 µM actin cross-linked by 20 mol% 20 nm low affinity cross-linker. ii. Representative image of an actin network (red) with 20 mol% 20 nm low affinity cross-linker 7 min 30 s after adding 0.24 µM myosin II (white). (C) Mean distance between myosin clusters over time in entangled networks (gray), networks with 2.5 mol% 20 nm low affinity cross-linker (red) and 20 mol% 20 nm low affinity cross- linker (blue). Shaded regions indicate the window of time used to track the myosin cluster merges in Fig. 3F. Scale bar is 10 µm and time is represented in min:s in all images.

Although the low affinity cross-linker does not visibly bundle the actin filaments, we investigated myosin II-induced contractility to test the connectivity in the networks. When we introduce myosin II motors to a 2.5 mol% cross-linked network, we observe bending fluctuations in the actin filaments as myosin incorporate in the network and form initial clusters. Then, as myosin clusters grow, the actin filaments straighten, indicating a cross-linked network (Fig. 2A ii). Once the dense regions form, the network begins to move collectively, consistent with the contraction signature in the high affinity cross-linker. During this period, the myosin clusters move closer together, with some merging with nearby clusters, which is consistent with the contractile cross-linked networks. At longer times, however, the myosin clusters merge into isolated, large actomyosin clusters, much like entangled actin networks. By contrast, when we increase the low affinity cross-linker concentration to 20 mol%, we see regions of myosin puncta forming within the network and buckling. The network continues to slowly evolve, as actin continues to buckle and flow, but myosin clustering is significantly slower (Fig. 2B ii), suggesting the network is approaching a frozen steady state.

The mean distance between myosin clusters in the 2.5 mol% low affinity cross-linked network sharply decreases over the first 40 s, as myosin clusters accumulate in the network, then increases as myosin clusters merge (Fig. 2C, red). While the timescale is faster, the trend in myosin clustering is similar to the entangled network (Fig. 2C, gray, same data as Fig. 1D). The myosin clusters merge an average of 1.6 ± 0.9 times (n=30, Fig. 3F, green), similar to the high affinity cross-linker. By contrast, in the 20 mol% cross-linked networks, the mean distance between clusters remains stable over time (Fig. 2C, blue), like the high affinity cross-linker (Fig 1D, blue). Myosin clusters merge 0.4 ± 0.6 merges (n=39) in these networks (Fig. 3F, black), consistent with frozen network.

**Figure 3:**
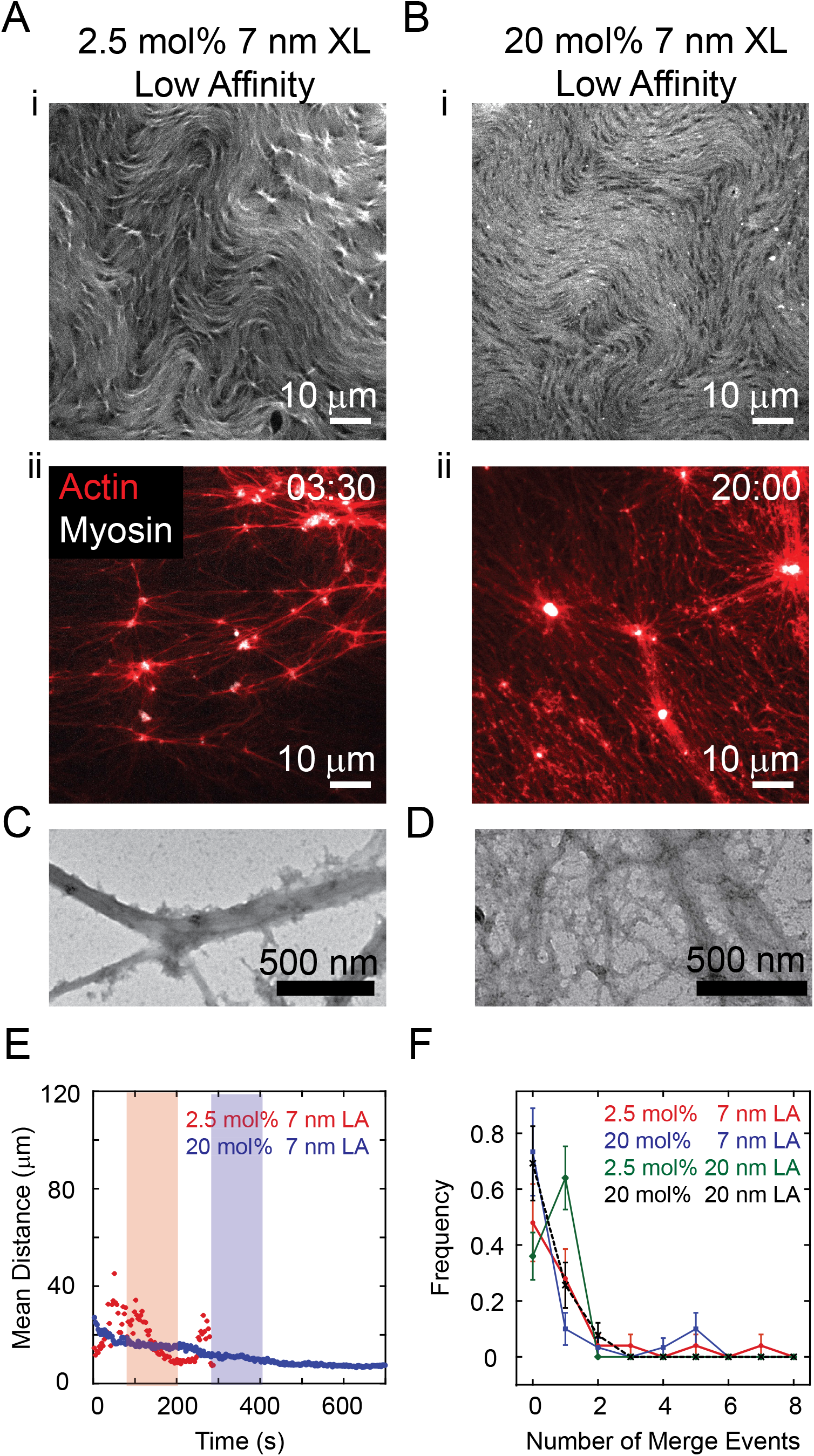
Short low affinity cross-linker forms contractile networks. (A) i. Fluorescence microscopy image of actin network cross-linked by 2.5 mol% 7 nm low affinity cross-linker and (ii.) 3 min 30 s after adding 0.12 µM myosin II (white, actin is in red). (B) i. Fluorescence microscopy image of actin network cross-linked by 20 mol% 7 nm low affinity cross-linker and (ii.) 20 min after 0.24 µM myosin II (white, actin is in red) was added. (C) Transmission electron microscopy image of actin filaments bundled by 20 mol% 7 nm low affinity cross-linker. (D) Transmission electron microscopy image of actin network cross-linked by 20 mol% 20 nm low affinity cross-linker. (E) Mean distance between myosin clusters for networks cross-linked by 7 nm low affinity cross-linker (red: 2.5 mol%, blue: 20 mol%). Shaded regions indicate the window of time used to track myosin merges in Fig. 3F. (F) Frequency of the number of times a myosin cluster merges for actin networks cross-linked by 20 nm low affinity cross-linker (green: 2.5 mol% and black: 20 mol%) and 7 nm low affinity cross-linker (red: 2.5 mol% and blue:20 mol%). Scale bar is 10 µm for fluorescence images and 500 nm for transmission electron microscopy images. Time is shown in min:s for all images.

### Short, low affinity cross-linkers form networks

In addition to binding affinity, cross-linker length is also known to influence the actin network microstructure^22^. Short physiological cross-linkers, with lower binding affinities than phalloidin, such as fascin and fimbrin, are known to bundle actin filaments^14,58^. Longer cross-linkers, like α- actinin also form networks and bundles, while long, flexible cross-linkers such as filamin are known to form fine mesh networks rather than bundles^59,18^. To test if shorter, but still low affinity cross-linkers would bundle actin, we synthesized a shorter linker using 20 base pairs of DNA, which is ∼7 nm long (compared to the long cross-linker, which is 60 base pairs or ∼20 nm long), similar to the physiological cross-linker, fascin^60^. We find that over a range of concentrations (2.5- 20 mol%), the 7 nm, low affinity cross-linker does not induce dominant bundling in the entangled actin network but may induce local regions to have higher densities of filaments (Fig. 3A i and Fig. 3B i). To further investigate the actin assembly microstructure, we imaged the cross-linked networks using transmission electron microscopy. At this higher magnification, we observe some bundled structures in the 7 nm, low affinity cross-linked samples (Fig. 3C), while the 20 nm, low affinity cross-linker forms a mesh network of filaments (Fig. 3D). This indicates that even though in fluorescence images the cross-linker length does not appear to influence the actin microstructure, the underlying microstructures between the two cross-linked networks differ.

Myosin II motors induce network remodeling in the short, low affinity cross-linked networks in the same qualitative manner as in the long, low affinity cross-linked networks. At 2.5 mol% cross- linker, the myosin accumulates quickly, concurrent with actin forming a straightened network of bundles between clusters. Then, as myosin clusters merge, the network undergoes mass contraction (Fig. 3A ii). At higher cross-linker concentration (20 mol%), as the myosin motors accumulate, they induce local filament bending and aggregation in the short, low affinity cross- linked network. The myosin clusters merge and grow as they slowly remodel the network, consistent with the longer low affinity cross-linkers (Fig. 3B ii).

When we quantify the mean distance between clusters, we also find similar trends in the low affinity cross-linkers at 2.5 mol% cross-linking (short, Fig. 3E, red and long, Fig. 2C, red) and 20 mol% cross-linking (short, Fig. 3E blue, and long, Fig. 2C, blue). The myosin clustering dynamics are similar across all cross-linked networks (Fig. 1E blue and Fig. 3F). We find that in the 2.5 mol% 7 nm low affinity cross-linked network, myosin cluster merge an average of 1.2 ± 1.6 times (n=25), while in 20 mol% networks, the average number of merges per cluster is 0.8 ± 1.6 times (n=30).

### Short, high affinity cross-linker forms a network of bundles

For completion, we also prepared networks using short, high affinity cross-linkers, even though the long, high affinity cross-linkers already exhibit actin bundling. To synthesize these cross- linkers, we used the same DNA sequence as for the short, low affinity cross-linkers, but with the biotin-streptavidin binding regions. In networks with 2.5 mol% of the short, high affinity cross- linker, we observe the rapid formation of bundles (Fig. 4A i). These bundles have an average diameter of 0.74 ± 0.21 µm (n=40 bundles), which is on the same order, but slightly larger than the bundles formed by the 20 nm high affinity cross-linker.

**Figure 4:**
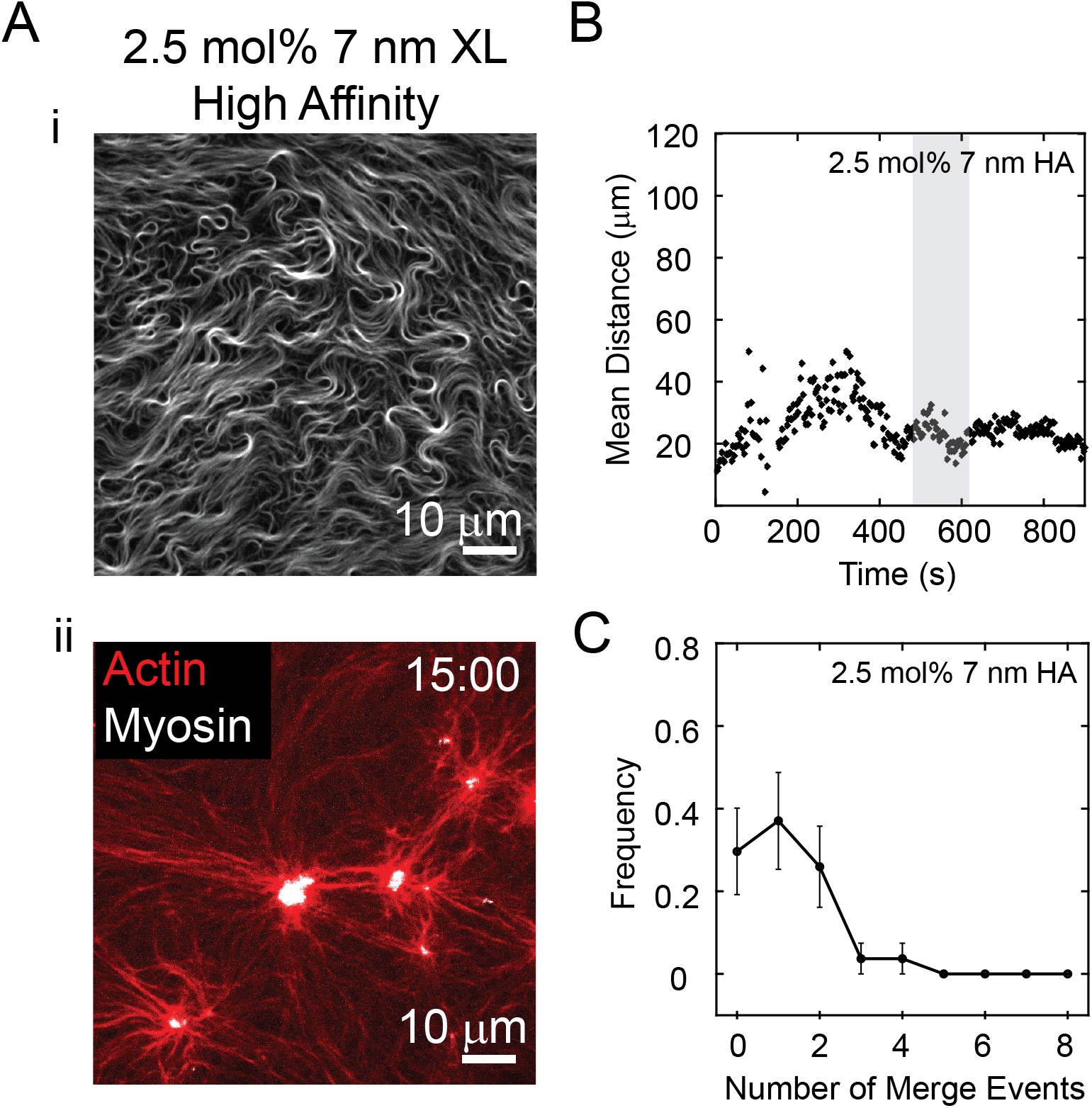
Short, high affinity cross-linker forms bundles. (A) i. Fluorescence microscopy image of actin network cross-linked by 2.5 mol% 7 nm high-binding affinity cross-linker and (ii.) 15 min after adding 0.12 µM myosin II (white, actin is in red). (B) Mean distance between myosin clusters within an actin network cross-linked by 2.5 mol% 7 nm high affinity cross-linker. (C) Frequency of the times a myosin cluster merges in actin networks cross-linked by 2.5 mol% 7 nm high affinity. The shaded region in Fig. 4B indicates the window of time used to track myosin cluster merges. Scale bar for all images is 10 µm and time is shown in min:s.

Myosin motors induce qualitatively similar activity in short, high affinity cross-linked networks as long, high affinity cross-linked networks (Fig. 4A ii). As myosin accumulates, actin filament bending occurs and the filaments arrange into an interconnected network of bundles between small clusters. As the myosin clusters continue to grow, the actin filaments become straighter, suggesting they are under tension, and begin to move as a collective network. As with the long, high affinity cross-linker, the mean distance between myosin clusters has a small maxima as the myosin accumulates in the network and initially merge, then decreases as larger myosin clusters move towards each other in the connected network, and then finally asymptotes as the network stabilizes (Fig. 4B, black). The myosin clusters merge an average of 1.2 ± 1.0 times (Fig. 4C, n=26).

## Conclusions

Here, we synthesized a programmable cross-linker, where cross-linker properties can be systematically modified to investigate the relationship between actin assembly microstructure, myosin II induced contraction, and the underlying cross-linking properties. We find that high affinity cross-linkers bundle actin filaments, while low affinity cross-linkers with the same lengths form cross-linked actin networks. Transmission electron microscopy reveals that low affinity cross-linkers can have mixed bundle and network microstructures at high cross-linking concentrations. Interestingly, myosin II remodels and contracts the cross-linked networks, for all lengths (7 nm and 20 nm) and affinities (K_D_∼10^-13^ M and K_D_∼10^-8^M), and concentrations (2.5 mol% and 20 mol%) tested, qualitatively differently than uncross-linked networks of entangled actin filaments. At the highest cross-linker concentrations (20 mol%), we find that myosin slowly remodels the network and approaches a frozen state. At low cross-linker concentrations (2.5 mol%), myosin driven network contraction occurs qualitatively similarly in all cases. However, as actomyosin clusters increase in size, the high affinity crosslinked networks retain connectivity, while in the low affinity cross-linked networks, disparate clusters merge while losing connectivity. This research expands our understanding of the structure-property relationship in subcellular materials, as well as soft materials design.

## Acknowledgements

This work was supported in part by the National Science Foundation EPSCoR Program under NSF Award #OIA-1655740. We thank a Department of Defense SMART Fellowship to C.A.F. and Clemson University’s Biophysics REU under National Science Foundation Award #2349368 with funding from the DBI and the EPSCoR program. We also thank Clemson University’s Core Incentivized Access award for electron microscopy use, as well as Donald Mulwee and Dr. Destiny Williams for their help.

## References

(1) Fletcher, D. A.; Mullins, R. D. Cell Mechanics and the Cytoskeleton. Nature 2010, 463 (7280), 485–492. 10.1038/nature08908

(2) Pollard, T. D. Actin and Actin-Binding Proteins. Cold Spring Harb Perspect Biol 2016, 8 (8), a018226. 10.1101/cshperspect.a018226

(3) Thoresen, T.; Lenz, M.; Gardel, M. L. Reconstitution of Contractile Actomyosin Bundles. Biophysical Journal 2011, 100 (11), 2698–2705. 10.1016/j.bpj.2011.04.031

(4) Kools, W.; Faour, S.; Carlier, A.; Hristova, T.; Sirkia, M. E.; Chaix, L.; Lee, A.; Martin, P.; Houdusse, A.; Romet-Lemonne, G.; Jégou, A. Myosin-X Sorts Actin Filaments into Parallel Bundles and Tunes Barbed-End Dynamics. Biochemistry September 11, 2026. 10.64898/2026.09.08.749704

(5) Graham, K.; Chandrasekaran, A.; Wang, L.; Ladak, A.; Lafer, E. M.; Rangamani, P.; Stachowiak, J. C. Liquid-like VASP Condensates Drive Actin Polymerization and Dynamic Bundling. Nat. Phys. 2023, 19 (4), 574–585. 10.1038/s41567-022-01924-1

(6) Feigeles, C. A.; Brasovs, A.; Puchalski, A.; Laukat, O.; Kornev, K. G.; Weirich, K. L. Capillary-Induced Bundling of Biopolymer Networks via Protein Condensates. January 25, 2025. 10.1101/2025.01.22.634359

(7) Gouveia, B.; De Souza, J. P.; Valdez, V.; Shaevitz, J. W.; Stone, H. A.; Petry, S. Capillary Bundling of Microtubules by Condensates. Biophysics June 20, 2026. 10.64898/2026.06.19.733462

(8) Wiegand, T.; Hyman, A. A. Drops and Fibers — How Biomolecular Condensates and Cytoskeletal Filaments Influence Each Other. Emerging Topics in Life Sciences 2020, 4 (3), 247–261. 10.1042/ETLS20190174

(9) Sanchez, T.; Chen, D. T. N.; DeCamp, S. J.; Heymann, M.; Dogic, Z. Spontaneous Motion in Hierarchically Assembled Active Matter. Nature 2012, 491 (7424), 431–434. 10.1038/nature11591

(10) Conway, L.; Gramlich, M. W.; Ali Tabei, S. M.; Ross, J. L. Microtubule Orientation and Spacing within Bundles Is Critical for Long-range Kinesin-1 Motility. Cytoskeleton 2014, 71 (11), 595–610. 10.1002/cm.21197

(11) Tharmann, R.; Claessens, M. M. A. E.; Bausch, A. R. Micro- and Macrorheological Properties of Actin Networks Effectively Cross-Linked by Depletion Forces. Biophysical Journal 2006, 90 (7), 2622–2627. 10.1529/biophysj.105.070458

(12) Blanchoin, L.; Boujemaa-Paterski, R.; Sykes, C.; Plastino, J. Actin Dynamics, Architecture, and Mechanics in Cell Motility. Physiological Reviews 2014, 94 (1), 235–263. 10.1152/physrev.00018.2013

(13) Vignjevic, D.; Kojima, S.; Aratyn, Y.; Danciu, O.; Svitkina, T.; Borisy, G. G. Role of Fascin in Filopodial Protrusion. The Journal of Cell Biology 2006, 174 (6), 863–875. 10.1083/jcb.200603013

(14) Edwards, R. A.; Bryan, J. Fascins, a Family of Actin Bundling Proteins. Cell Motil. Cytoskeleton 1995, 32 (1), 1–9. 10.1002/cm.970320102

(15) Mogilner, A.; Rubinstein, B. The Physics of Filopodial Protrusion. Biophysical Journal 2005, 89 (2), 782–795. 10.1529/biophysj.104.056515

(16) Li, T.; Oloyede, A.; Gu, Y. T. F-Actin Crosslinker: A Key Player for the Mechanical Stability of Filopodial Protrusion. Journal of Applied Physics 2013, 114 (21), 214701. 10.1063/1.4839715

(17) Schmoller, K. M.; Lieleg, O.; Bausch, A. R. Structural and Viscoelastic Properties of Actin/Filamin Networks: Cross-Linked versus Bundled Networks. Biophysical Journal 2009, 97 (1), 83–89. 10.1016/j.bpj.2009.04.040

(18) Tseng, Y.; An, K. M.; Esue, O.; Wirtz, D. The Bimodal Role of Filamin in Controlling the Architecture and Mechanics of F-Actin Networks. Journal of Biological Chemistry 2004, 279 (3), 1819–1826. 10.1074/jbc.M306090200

(19) Noureddine, M.; Mikolajek, H.; Morgan, N. V.; Denning, C.; Loughna, S.; Gehmlich, K.; Mohammed, F. Structural and Functional Insights into α-Actinin Isoforms and Their Implications in Cardiovascular Disease. Journal of General Physiology 2025, 157 (2), e202413684. 10.1085/jgp.202413684

(20) Pecorari, I.; Mestroni, L.; Sbaizero, O. Current Understanding of the Role of Cytoskeletal Cross-Linkers in the Onset and Development of Cardiomyopathies. IJMS 2020, 21 (16), 5865. 10.3390/ijms21165865

(21) Zhang, L.; Zhang, N.; Tang, X.; Liu, F.; Luo, S.; Xiao, H. Increased α-Actinin-2 Expression in the Atrial Myocardium of Patients with Atrial Fibrillation Related to Rheumatic Heart Disease. Cardiology 2016, 135 (3), 151–159. 10.1159/000446362

(22) Lieleg, O.; Claessens, M. M. A. E.; Bausch, A. R. Structure and Dynamics of Cross-Linked Actin Networks. Soft Matter 2010, 6 (2), 218–225. 10.1039/B912163N

(23) Daly, M. L.; Nishi, K.; Klawa, S. J.; Hinton, K. Y.; Gao, Y.; Freeman, R. Designer Peptide–DNA Cytoskeletons Regulate the Function of Synthetic Cells. Nat. Chem. 2024, 16 (8), 1229–1239. 10.1038/s41557-024-01509-w

(24) Lorenz, J. S.; Schnauß, J.; Glaser, M.; Sajfutdinow, M.; Schuldt, C.; Käs, J. A.; Smith, D. M. Synthetic Transient Crosslinks Program the Mechanics of Soft, Biopolymer-Based Materials. Advanced Materials 2018, 30 (13), 1706092. 10.1002/adma.201706092

(25) Jorgenson, T. D.; Baboolall, K. D.; Suarez, C.; Kovar, D. R.; Gardel, M. L.; Rowan, S. J. Highly Flexible PEG-LifeAct Constructs Act as Tunable Biomimetic Actin Crosslinkers. Soft Matter 2024, 20 (5), 971–977. 10.1039/D3SM01341C

(26) Nagy, S.; Ricca, B. L.; Norstrom, M. F.; Courson, D. S.; Brawley, C. M.; Smithback, P. A.; Rock, R. S. A Myosin Motor That Selects Bundled Actin for Motility. Proc. Natl. Acad. Sci. U.S.A. 2008, 105 (28), 9616–9620. 10.1073/pnas.0802592105

(27) Scholz, M.; Burov, S.; Weirich, K. L.; Scholz, B. J.; Tabei, S. M. A.; Gardel, M. L.; Dinner, A. R. Cycling State That Can Lead to Glassy Dynamics in Intracellular Transport. Phys. Rev. X 2016, 6 (1), 011037. 10.1103/PhysRevX.6.011037

(28) Ali, M. Y.; Previs, S. B.; Trybus, K. M.; Sweeney, H. L.; Warshaw, D. M. Myosin VI Has a One Track Mind Versus Myosin VA When Moving on Actin Bundles or at an Intersection. Traffic 2013, 14 (1), 70–81. 10.1111/tra.12017

(29) Weirich, K. L.; Stam, S.; Munro, E.; Gardel, M. L. Actin Bundle Architecture and Mechanics Regulate Myosin II Force Generation. Biophysical Journal 2021, 120 (10), 1957–1970. 10.1016/j.bpj.2021.03.026

(30) Murrell, M. P.; Gardel, M. L. F-Actin Buckling Coordinates Contractility and Severing in a Biomimetic Actomyosin Cortex. Proc. Natl. Acad. Sci. U.S.A. 2012, 109 (51), 20820–20825. 10.1073/pnas.1214753109

(31) Reymann, A.-C.; Boujemaa-Paterski, R.; Martiel, J.-L.; Guérin, C.; Cao, W.; Chin, H. F.; De La Cruz, E. M.; Théry, M.; Blanchoin, L. Actin Network Architecture Can Determine Myosin Motor Activity. Science 2012, 336 (6086), 1310–1314. 10.1126/science.1221708

(32) Stam, S.; Freedman, S. L.; Banerjee, S.; Weirich, K. L.; Dinner, A. R.; Gardel, M. L. Filament Rigidity and Connectivity Tune the Deformation Modes of Active Biopolymer Networks. Proc. Natl. Acad. Sci. U.S.A. 2017, 114 (47). 10.1073/pnas.1708625114

(33) Koenderink, G. H.; Paluch, E. K. Architecture Shapes Contractility in Actomyosin Networks. Current Opinion in Cell Biology 2018, 50, 79–85. 10.1016/j.ceb.2018.01.015

(34) Muresan, C. G.; Sun, Z. G.; Yadav, V.; Tabatabai, A. P.; Lanier, L.; Kim, J. H.; Kim, T.; Murrell, M. P. F-Actin Architecture Determines Constraints on Myosin Thick Filament Motion. Nat Commun 2022, 13 (1), 7008. 10.1038/s41467-022-34715-6

(35) Ennomani, H.; Letort, G.; Guérin, C.; Martiel, J.-L.; Cao, W.; Nédélec, F.; De La Cruz, E. M.; Théry, M.; Blanchoin, L. Architecture and Connectivity Govern Actin Network Contractility. Current Biology 2016, 26 (5), 616–626. 10.1016/j.cub.2015.12.069

(36) Bendix, P. M.; Koenderink, G. H.; Cuvelier, D.; Dogic, Z.; Koeleman, B. N.; Brieher, W. M.; Field, C. M.; Mahadevan, L.; Weitz, D. A. A Quantitative Analysis of Contractility in Active Cytoskeletal Protein Networks. Biophysical Journal 2008, 94 (8), 3126–3136. 10.1529/biophysj.107.117960

(37) Stam, S.; Gardel, M. L.; Weirich, K. L. Direct Detection of Deformation Modes on Varying Length Scales in Active Biopolymer Networks. Biophysics May 16, 2023. 10.1101/2023.05.15.540780

(38) Stam, S.; Huntley, S.; Feigeles, C. A.; Armstrong, V. J.; Cheves, M. A.; Rubin, S.; Weirich, K. L. Tuning the Contractility and Deformation Modes of Active Actin-Based Assemblies in Vitro: From 2D Active Networks to Liquid Crystal Drops. December 8, 2024. 10.1101/2024.12.04.626882

(39) Spudich, J. A.; Watt, S. The Regulation of Rabbit Skeletal Muscle Contraction. Journal of Biological Chemistry 1971, 246 (15), 4866–4871. 10.1016/S0021-9258(18)62016-2

(40) Verkhovsky, A. B.; Borisy, G. G. Non-Sarcomeric Mode of Myosin II Organization in the Fibroblast Lamellum. The Journal of cell biology 1993, 123 (3), 637–652. 10.1083/jcb.123.3.637

41. Weber, P. C.; Ohlendorp, D. H.; Wendolosiu, J. J. Structural Origins of High-Affinity Biotin Binding to Streptavidin. 243.

(42) Chilkoti, A.; Stayton, P. S. Molecular Origins of the Slow Streptavidin-Biotin Dissociation Kinetics. J. Am. Chem. Soc. 1995, 117 (43), 10622–10628. 10.1021/ja00148a003

(43) Dundas, C. M.; Demonte, D.; Park, S. Streptavidin–Biotin Technology: Improvements and Innovations in Chemical and Biological Applications. Appl Microbiol Biotechnol 2013, 97 (21), 9343–9353. 10.1007/s00253-013-5232-z

(44) Holden, B.; Mutaher, M.; Raychowdhury, R.; Bakalar, M.; Lopez, A.; Schwartz, M. A.; Olajide, E.; Santos, M.; Hacohen, N. An Affinity Reagent-Conjugated Biotin Ligase for Amplified Cell Surface Labelling in Vitro and in Vivo. Sci Rep 2026, 16 (1), 18463. 10.1038/s41598-026-46452-7

(45) Khatri, S.; He, J. Probing Streptavidin-Biotin Interaction and Complex Structure at the Single-Molecule Level Using a Self-Crowded Nanopipette. Small 2026, 22 (11), e08579. 10.1002/smll.202508579

(46) Pollard, T. D.; Cooper, J. A. Actin, a Central Player in Cell Shape and Movement. Science 2009, 326 (5957), 1208–1212. 10.1126/science.1175862

(47) Lee, D. T.; Schachter, B. J. Two Algorithms for Constructing a Delaunay Triangulation. International Journal of Computer and Information Sciences 1980, 9 (3), 219–242. 10.1007/BF00977785

(48) Musin, O. R. Properties of the Delaunay Triangulation. In Proceedings of the thirteenth annual symposium on Computational geometry - SCG ’97; ACM Press: Nice, France, 1997; pp 424–426. 10.1145/262839.263061

(49) Belmonte, J. M.; Leptin, M.; Nédélec, F. A Theory That Predicts Behaviors of Disordered Cytoskeletal Networks. Mol Syst Biol 2017, 13 (9), MSB177796. 10.15252/msb.20177796

(50) Wachsstock, D. H.; Schwartz, W. H.; Pollard, T. D. Affinity of Alpha-Actinin for Actin Determines the Structure and Mechanical Properties of Actin Filament Gels. Biophysical Journal 1993, 65 (1), 205–214. 10.1016/S0006-3495(93)81059-2

(51) Ohta, Y.; Hartwig, J. H. Actin Filament Crosslinking by Chicken Gizzard Filamin Is Regulated by Phosphorylation in Vitro. Biochemistry 1995, 34 (20), 6745–6754. 10.1021/bi00020a020

(52) Xu, J.; Wirtz, D.; Pollard, T. D. Dynamic Cross-Linking by α-Actinin Determines the Mechanical Properties of Actin Filament Networks. Journal of Biological Chemistry 1998, 273 (16), 9570–9576. 10.1074/jbc.273.16.9570

(53) Chaubet, L.; Chaudhary, A. R.; Heris, H. K.; Ehrlicher, A. J.; Hendricks, A. G. Dynamic Actin Cross-Linking Governs the Cytoplasm’s Transition to Fluid-like Behavior. MBoC 2020, 31 (16), 1744–1752. 10.1091/mbc.E19-09-0504

(54) Lengsfeld, A. M.; Lowt, I.; Wielandt, T.; Dancker, P.; Hasselbach, W. Interaction of Phalloidin with Actin. Proc. Nat. Acad. Sci. USA 1974.

(55) Faulstich, H.; Schäfer, A. J.; Weckauf, M. The Dissociation of the Phalloidin-Actin Complex. Hoppe-Seyleŕs Zeitschrift für physiologische Chemie 1977, 358 (1), 181–184. 10.1515/bchm2.1977.358.1.181

(56) Green, N. M. Avidin. In Advances in Protein Chemistry; Elsevier, 1975; Vol. 29, pp 85–133. 10.1016/S0065-3233(08)60411-8

(57) Visegrády, B.; Lőrinczy, D.; Hild, G.; Somogyi, B.; Nyitrai, M. The Effect of Phalloidin and Jasplakinolide on the Flexibility and Thermal Stability of Actin Filaments. FEBS Letters 2004, 565 (1–3), 163–166. 10.1016/j.febslet.2004.03.096

(58) Bretscher, A. Fimbrin Is a Cytoskeletal Protein That Crosslinks F-Actin in Vitro. Proc. Natl. Acad. Sci. U.S.A. 1981, 78 (11), 6849–6853. 10.1073/pnas.78.11.6849

(59) Pelletier, O.; Pokidysheva, E.; Hirst, L. S.; Bouxsein, N.; Li, Y.; Safinya, C. R. Structure of Actin Cross-Linked with α -Actinin: A Network of Bundles. Phys. Rev. Lett. 2003, 91 (14), 148102. 10.1103/PhysRevLett.91.148102

(60) Kureishy, N.; Sapountzi, V.; Prag, S.; Anilkumar, N.; Adams, J. C. Fascins, and Their Roles in Cell Structure and Function. BioEssays 2002, 24 (4), 350–361. 10.1002/bies.10070

