## Supporting Information for "Cross-linker binding strength influences actin assembly microstructure and myosin II driven remodeling"

**Biotinylated Sample Preparation**

To use biotin-streptavidin as a binding interaction, we first needed to determine the concentration of biotinylated actin that would not cause bundling with free streptavidin in solution. We chose to work at 5 mol% biotinylated actin and 2.5 mol% streptavidin for biotinylated cross-linker samples because of the negligible streptavidin-induced bundling.

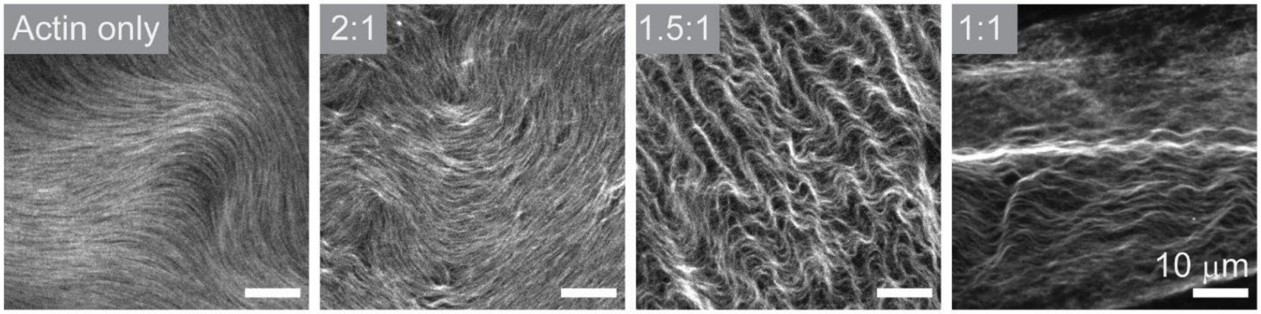


**Fig. S1. Determination of biotin:streptavidin molar ratio that does not induce bundle formation.** Fluorescence microscopy images of (A) biotinylated actin filaments forming an entangled network (2.64 μM actin, 5 mol% biotinylated). Fluorescence microscopy images of biotinylated actin networks with (B) 2.5 mol% streptavidin (2:1 biotin:streptavidin) (C) 3.3 mol% (1.5:1 biotin:streptavidin), and (D) 5 mol% (1:1 biotin:streptavidin). Scale bar is 10 μm in all images.

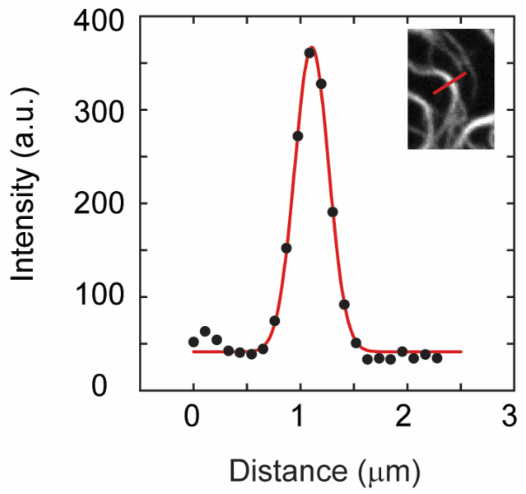


**Fig. S2.** Representative data used to determine bundle thickness. The intensity (black, circles) along a line perpendicular to the length of an actin bundle (inset, red line) with Gaussian fit (red).

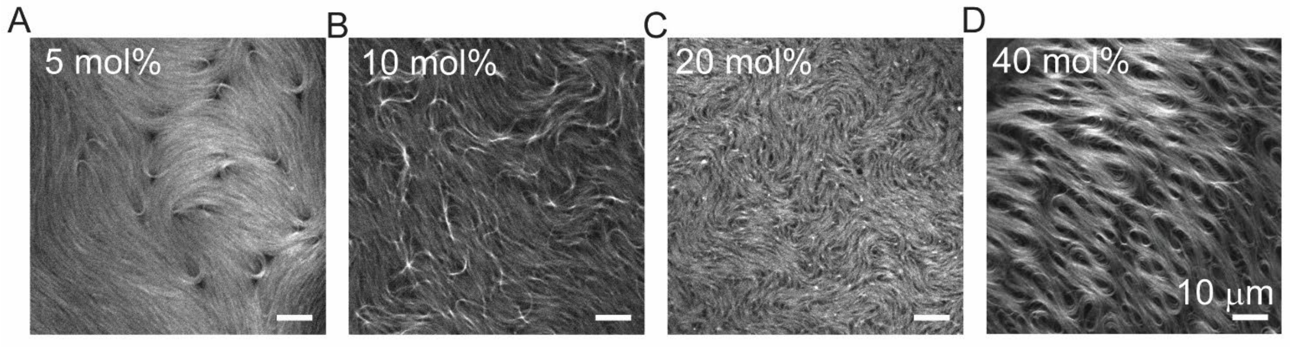


**Fig. S3.** (A) Fluorescence microscopy images of 2.64 μM actin cross-linked by (A) 5 mol%, (B) 10 mol%, (C) 20 mol%, and (D) 40 mol% 20 nm low affinity cross-linker. Scale bar is 10 μm.
